# Highly structured habitats mitigate size- and growth-selective mortality of post-settlement juvenile fish

**DOI:** 10.1101/2022.10.21.513133

**Authors:** Yasuhiro Kamimura, Jun Shoji

**Affiliations:** Takehara Fisheries Research Station, Graduate School of Biosphere Science, Hiroshima University, 5-8-1 Minato-Machi, Takehara, Hiroshima 725-0024, Japan

**Author notes:** **Corresponding author:** Y. Kamimura. Fisheries Resource Institute, Japan Fisheries Research and Education Agency, 2-12-4 Fukuura, Kanazawa, Yokohama, Kanagawa 236-8648, Japan (Yasuhiro Kamimura). Division of Marine Science and Technology, Fukui Prefectural University, 1-1 Gakuen-cho, Obama, Fukui 917-0013, Japan (Jun Shoji).

**Keywords:** Growth-selective predation, size-selective predation, temperature, macroalgal bed, *Sargassum*, *Sebastes cheni*, mortality rate coefficient

## Abstract

The role of vegetated habitats such as seagrass and macroalgal beds as nurseries is essential for the survival of larval and juvenile fish, although quantitative evaluation of the contribution of these habitats to nursery function is limited. Moreover, growth–survival relationships of larvae and juveniles associated with vegetated habitats have rarely been examined. To quantitatively and qualitatively evaluate the processes affecting juvenile survival in vegetated habitats, we investigated whether there is a correlation between the degree of selection for bigger and faster-growing fish and mortality rates for three cohorts by birth date of post-settlement black rockfish (*Sebastes cheni*) in a macroalgal bed. We also analyzed relationships between growth rate and experienced temperature by age class to examine the effects of temperature on growth. The latest cohort, which grew under lower vegetation coverage due to a seasonal increase in water temperature, showed higher mortality with evidence of strong selection for bigger and faster-growing fish. The growth–temperature relationships showed that positive effects of temperature on growth weakened after settlement. Therefore, we suggest that macroalgal coverage has a critical role controlling the growth–mortality relationship of post-settlement *S. cheni*. Furthermore, the negative effects of high temperature on juvenile survival through loss of vegetation may be greater than the positive effects on juvenile growth. These findings will contribute to the management of fisheries resources by increasing the understanding of relationships between survival mechanisms in fish early life stages and vegetation phenology of their habitat under the increasing effects of global warming.

## Introduction

Vegetated habitats such as seagrass and algal beds have long been recognized as highly productive areas and important nursery grounds for marine organisms (e.g., Heck et al.., 2003). Indeed, the contribution of these beds to food production is estimated to be among the most valuable ecosystem services in the world (Costanza et al., 2014); moreover, the value of these beds is likely to have been underestimated in previous analyses because the analyses relied on only a few studies (de Groot et al., 2012) that did not systematically evaluate the value of seagrass and algal beds around the globe (TEEB, 2012). In particular, the quantitative evaluation of the contribution of algal beds as nurseries to fish production is lacking compared to that of seagrass beds (Lefcheck et al., 2019; McDevitt-Irwin et al., 2016). Hence, further quantitative and qualitative studies are needed to examine the nursery role of seagrass and algal beds, i.e., the mechanisms behind variability in density, growth, and survival of larval and juvenile fishes.

For the population dynamics of fish early life stages, growth rates of cohorts and populations are important survival indices, with faster growth and higher survival leading to recruitment success (Anderson, 1988). This “growth–mortality” hypothesis includes three non-exclusive functional hypotheses. First, fast growth shortens the duration of larval stages when the proportion of mortality due to starvation, predation, and physical processes is highest (the “stage duration” hypothesis; Chambers & Leggett, 1987; Houde, 1987). Second, fast growth can increase the probability of survival for bigger and faster-growing individuals because they are less vulnerable to predation compared to smaller and slower-growing individuals (the “bigger-is-better” hypothesis; Miller et al., 1988). Third, growth rate itself also contributes to the probability of survival for fish in early life stages regardless of fish size (the “growth-selective predation” hypothesis; Takasuka et al., 2003), as it reflects antipredator behavior such as burst swimming speed (Nakamura et al., 2022). Many studies have tested these hypotheses in a variety of fishes and ecosystems (e.g., Fennie et al., 2020; Hare & Cowen, 1997; Islam et al., 2010; Takasuka et al., 2017). Moreover, these growth-dependent mortality processes extend beyond the larval stage into juvenile stages (Khamassi et al., 2020; Sogard, 1997).

Previous field studies have reported positive selection for fast-growing individuals. The survivors (SV) of a cohort of fish sampled later in the early life stages had faster growth rates than those sampled earlier (original population, OP) during the same period, showing that faster-growing individuals had a better chance for survival (e.g., Meekan & Fortier, 1996; Shoji & Tanaka, 2006; Takasuka et al., 2017). However, there are few studies that directly determine how the degree of selection for growth (positive or negative) correlates with the mortality rate of the cohort (e.g., Holmes & McCormick, 2006; Takasuka et al., 2007). To achieve that, we need sampling designs that enable quantitative analysis of mortality rates together with the degree of size and growth selection within and among cohorts in field surveys, laboratory experiments, or mesocosm studies.

In coastal ecosystems, habitat complexity, such as seagrass shoot density or vegetation coverage in algal beds, is likely to be an important determinant of larval and juvenile mortality in both mobile and sessile animals (e.g., Heck & Orth, 2006; Johnson, 2006) because it determines the effective shelter for prey against transient predators (Horinouchi, 2007). Moreover, habitat characteristics likely modify or mask the effects of size-selective mortality in early life stages of teleost fishes (Sogard, 1997), although that has not been directly demonstrated. It would be helpful to investigate any linkage between mortality rate and size- and growth-selective mortality in habitats with different complexity to understand the nursery function of vegetated habitats more comprehensively.

Black rockfish (*Sebastes cheni*) is a commercially important fishery resource in temperate coastal waters around Japan and the southern part of the Korean Peninsula (Kai & Nakabo, 2008). Juveniles settle in vegetated habitats such as seagrass (*Zostera marina*) and macroalgal (*Sargassum* spp.) beds at about 20 mm total length (TL) in spring and grow to about 60 mm by late May (Kamimura et al., 2011). Survival during such substrate-associated periods for the genus *Sebastes* is probably related to recruitment success (Laidig et al., 2007; Ralston et al., 2013). The effect of vegetation coverage on the cohort-specific mortality rate of juvenile *S. cheni* was previously studied in a macroalgal bed (mainly temperate *Sargassum* spp.) by sampling the same cohort repeatedly at relatively short time intervals (1–2 weeks) (Kamimura & Shoji, 2013). The later cohorts had higher mortality rates during the post-settlement period due to a decrease in vegetation coverage, which is considered a refuge for juveniles from nighttime predators (Kinoshita et al., 2014). In addition, seasonal increases in water temperature caused both a decrease in vegetation cover and higher growth rates in the later cohorts (Kamimura & Shoji, 2013). However, it is still not clear how water temperature affected *S. cheni* growth at the individual level, or how it related to growth–survival mechanisms in the macroalgal bed.

This study focused on the quantitative and qualitative understanding of the survival mechanisms of post-settlement *S. cheni* juveniles to help evaluate the fish nursery function of a macroalgal bed. First, to determine the growth–survival relationship of post-settlement *S. cheni*, we compared growth trajectories between individuals sampled in earlier and those in later using three cohorts based on birth date that reportedly experienced different mortality rates in a previous study (Kamimura & Shoji, 2013). We then examined the linkage between the degree of selection for bigger and faster-growing fish and mortality in cohorts of post-settlement juvenile *S. cheni*. We also investigated how growth trajectories differed among cohorts and how water temperature affected growth by estimating the temperatures experienced by individuals from birth to capture. Finally, we discuss the environmental conditions (vegetation coverage in the macroalgal bed and water temperature) that affect growth and survival of *S. cheni*.

## Materials and methods

### Field sampling

Field surveys were conducted in coastal waters off Abajima Island in Takehara, Hiroshima, on the central Seto Inland Sea, Japan (Figure 1) from February to May 2008. Information about the sampling site is detailed elsewhere (Kamimura & Shoji, 2013). Because the vegetation is dominated by patches of seagrass *Zostera marina* during summer and macroalgae (mostly *Sargassum* spp.) from winter to late spring (Kamimura & Shoji, 2009), we regarded this site as a macroalgal bed from February to May. Biotic and abiotic surveys were carried out during daytime (0900–1700 h local time) with a tidal level of 50–150 cm, in reference to mean lower low water, on each day at intervals of 5–14 days from 7 February to 30 May. Fish were collected using a round seine net (30 m long, 2 m high, 5-mm mesh aperture). In the macroalgal bed a square was enclosed on three sides using a net (10 m per side, 100 m^2^), with the fourth side facing the shoreline. All fish sampling was conducted at four separate areas randomly selected within the macroalgal bed.

**Figure 1.**
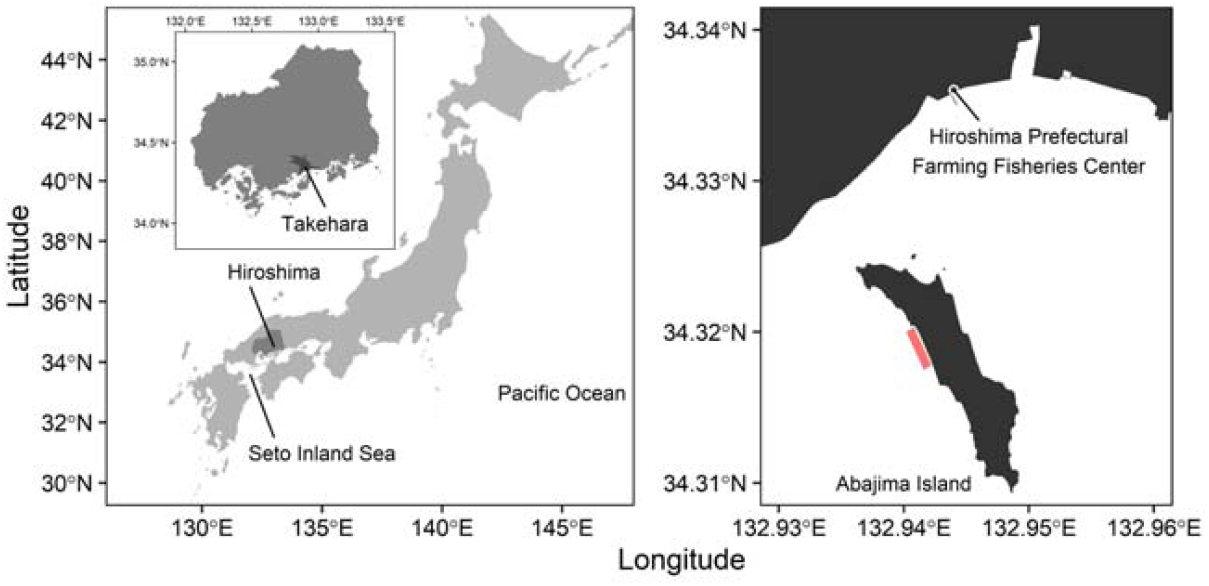
Location of Abajima Island in Takehara, Hiroshima, central Seto Inland Sea, Japan. The inset drawing in the left panel is an enlarged view of Hiroshima. The red shaded area in the right panel is the sampling site off the western coast of Abajima Island. Daily water temperature was obtained from the Hiroshima Prefectural Farming Fisheries Center, shown in the right panel.

Collected *Sebastes* juveniles were preserved in 10% formalin in seawater and subsamples were preserved in 90% ethanol for otolith analysis. Among the three *Sebastes* species—*S. inermis, S. ventricosus, and S. cheni*—*S. cheni* is commonly collected in the survey area (Kamimura et al., 2011). In the laboratory, *S. cheni* was identified on the basis of morphological differences (Kai & Nakabo, 2008) and measured for TL (mm) to the nearest 0.1 mm. The abundance of juvenile *S. cheni* on each sampling date is expressed as the number of fish per 100 square meters (inds. 100 m^−2^; Figure S1). Water temperatures in the macroalgal bed were measured using a multiple environmental measurement system (Alec Electronics Co., Ltd., Kobe, Japan). A vegetation coverage index (macroalgal occupancy in the water column: %) was estimated by visual census using a 1-m^2^ quadrat at four randomly selected separate areas. Seasonal changes in the juvenile abundance, and physical and biological environmental properties (water temperature, salinity, vegetation index, and zooplankton concentration) have been previously published (Kamimura & Shoji, 2013). Daily water temperature was obtained from the Hiroshima Prefectural Farming Fisheries Center (Figure 1) and used to estimate the temperature experienced by each individual fish (Figure S2).

### Mortality rate estimation

A total of 388 otoliths were processed in a previous study (Kamimura & Shoji, 2013) to determine birth dates to estimate cohort-specific mortality of juvenile *S. cheni*. In that study, juvenile *S. cheni* were divided into seven specific cohorts with 7-day periods based on individual birth dates. Cohorts were designated by alphabetical characters: A (25–31 Dec 2007), B (1–7 Jan 2008), C (8–14 Jan 2008), D (15–21 Jan 2008), E (22–28

Jan 2008), F (29 Jan–4 Feb 2008), and G (5–11 Feb 2008). Cohort A was excluded from further analysis because of its small sample size. Pairs of consecutive cohorts were then combined to make three new cohorts (I, II, and III), each of which included two of the original cohorts. Consequently, each new cohort spans two weeks of birth dates (Table 1 and Figure S3).

**Table 1.** Summary of cohort, birth and sampling dates (in 2008), sampling interval, number of individuals (*N*), mean (standard deviation, D) total length (TL), TL range, mean (SD) age, age range, and estimated mortality coefficient (*Z*) of juvenile *Sebastes cheni* from the central Seto Inland Sea, Japan, processed for growth analysis. Juveniles collected on the first date of the period used for mortality estimates for each cohort were regarded as the original population (OP), and those collected on subsequent dates were regarded as survivors (SV).

| Cohort | Birth date | Sampling date | Interval (days) | Group | *N* | TL (mm) |  | Age (days) |  | *Z* |
| --- | --- | --- | --- | --- | --- | --- | --- | --- | --- | --- |
|  |  |  |  |  |  | Mean (SD) | Range | Mean (SD) | Range |  |
| I | 1–14 Jan | 24 Mar | - | OP | 25 | 25.8 (1.4) | 23.7–29.1 | 76 (4) | 70–82 | 0.053 |
|  |  | 3 Apr | 10 | SV | 33 | 29.3 (2.2) | 25.4–35.9 | 85 (4) | 80–93 |  |
| II | 15–28 Jan | 3 Apr | - | OP | 21 | 27.0 (1.6) | 21.9–29.1 | 74 (4) | 67–79 | 0.044 |
|  |  | 16 Apr | 13 | SV | 14 | 32.0 (1.6) | 29.8–35.7 | 88 (3) | 83–92 |  |
| III | 29 Jan–11 Feb | 23 Apr | - | OP | 14 | 28.8 (1.7) | 25.9–31.0 | 80 (4) | 72–85 | 0.071 |
|  |  | 1 May | 8 | SV | 16 | 32.8 (2.1) | 30.3–37.1 | 87 (4) | 82–93 |  |

Instantaneous mortality coefficients (*Z*, day^−1^) of the three cohorts were estimated by applying an exponential model of decline (e.g., Secor & Houde, 1995) beginning after the day of maximum mean abundance of juveniles in the macroalgal bed (24 Mar for cohort I, 3 Apr for cohort II, and 23 Apr for cohort III):

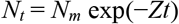

where *N*_*t*_ is the juvenile abundance at time *t* (days), *N*_*m*_ is the estimated abundance at the day of the maximum abundance for each cohort, and *Z* is the instantaneous daily mortality coefficient. Kamimura & Shoji (2013) considered the possible sampling bias due to net avoidance to be minimal because there was no day–night difference in the abundance of each size class. In addition, because a previous study reported no significant relationship between kelp density and emigration rates of juvenile *S. atrovirens* (Johnson, 2006), we assumed for the present study that emigration rates of *S. cheni* did not change with changes in vegetation coverage.

### Otolith analysis for growth estimation

Otolith daily increments were analyzed to estimate the birth dates and growth trajectories of juvenile *S. cheni*. The formation of daily otolith increments and extrusion check were validated by using cultured larvae and juveniles (Kamimura et al., 2012). The lapilli of a maximum of 53 juveniles for each sampling date were removed from the fish under a dissecting microscope. Each lapillus was embedded in enamel resin on a slide glass and dried, then ground with 2000–10,000 grid lapping films until the nucleus was clearly visible. Daily rings were counted from the extrusion check to the edge of the otolith and the radius of each daily ring was measured at 400–1000× magnification under a light microscope using the otolith daily ring measurement system (Ratoc System Engineering, Co., Ltd., Tokyo, Japan). Birth dates were estimated from the collection date and age of each fish.

### Estimation of juvenile S. cheni growth trajectories

Juvenile TL- and growth rate (GR)-at-age were back-calculated by using the biological intercept method (Campana, 1990). A linear model was fitted to the relationship between otolith radius and TL according to previous studies (Kamimura et al., 2012; Plaza et al., 2002). The length-at-age of *S. cheni* was calculated from the following equation:

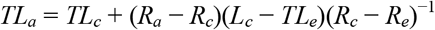

where *TL*_*a*_ and *TL*_*c*_ are fish size at age *a* and at capture, respectively; *R*_*a*_, *R*_*c*_, and *R*_*e*_ are otolith radius at age *a*, at capture, and at extrusion, respectively; and *TL*_*e*_ is mean fish size at extrusion (6.2 mm TL; Kamimura et al., 2012). GR-at-age was calculated from the difference between TL at age *a* + 1 and TL at age *a*. Because the outermost otolith increment may not have been fully formed, we excluded back-calculated TL and GR at that age or date (sampling date) from subsequent analyses.

### Intra-cohort comparison of growth trajectories

To examine the degree of selection for bigger and faster-growing individual *S. cheni* juveniles in the macroalgal bed, we compared the mean back-calculated TL and GR of each cohort earlier and later in the sampling date. Juveniles collected on the sampling date of maximum mean abundance for each cohort (24 March for cohort I, 3 April for cohort II, and 23 April for cohort III) (Table 1) were regarded as the original population (OP) for that cohort. Juveniles collected on the sampling date 8–13 days after the OP date (3 April for cohort I, 16 April for cohort II, and 1 May for cohort III) were regarded as survivors (SV) (Table 1). Back-calculated TL and GR were arranged by age and calendar date to compare growth trajectories between the two groups, OP and SV. The age-based comparison represents stage-specific differences, and the date-based comparison represents differences in growth characteristics (Takasuka et al., 2004). We therefore conducted both comparisons to examine stage-specific changes in growth trajectory, and post-settlement size- and growth-selective survival, using the 3-day mean back-calculated TL and GR. In the date-based comparison, the mean TL and the mean GR up to one month prior to the sampling date were compared between OP and SV by cohort.

### Inter-cohort comparison of GR and experienced temperature, and estimates of the growth–temperature relationship

Because growth rate is often affected by an individual’s age (Ashworth et al., 2017), we applied an appropriate model for the relationship between growth rate and age to remove any age effect on growth rate for the inter-cohort comparisons. Then, the model residuals were used to examine trends in growth and temperature effects on growth of *S. cheni*. We compared four models for the relationship between GR and age for *S. cheni*, following a previous study (Shima & Swearer, 2019), that is, a linear function, a quadratic function, a von Bertalanffy growth function, and a logistic growth function. A logistic growth model was selected as the appropriate model on the basis of Akaike’s information criterion (AIC), so the growth-rate residuals were used for the inter-cohort comparison and estimating the growth–temperature relationship (see Results). Experienced temperature for each fish was estimated from birth to collection using 3-day mean temperatures for this analysis.

### Statistical analysis

Instantaneous mortality coefficients were estimated by cohort using general linear models (LMs) with lognormal distribution. Repeated-measures multivariate analysis of variances (MANOVAs) were used for the age-based comparison of mean back-calculated TL between groups (OP and SV) by cohort. The mean back-calculated TLs for cohorts I, II, and III were compared up to 78 days, 72 days, and 72 days after birth, respectively, because of sample size. When conducting the date-based comparison, differences in birth date between groups (Table 1 and Figure S3) can affect the mean TL at a given calendar date. Therefore, multivariate analysis of covariances (MANCOVAs) were used with birth date (Julian date starting at 1 January 2008) as a covariate to compare mean back-calculated TLs between groups by cohort. Pillai’s Trace was selected as the statistic of MANOVAs and MANCOVAs because it is the most robust.

We used generalized linear models (GLMs) with gamma distribution and log link function to examine the differences in mean back-calculated GR-at-age between groups by cohort. Birth date was considered in the date-based GR comparison as with the date-based TL comparison. The mean back-calculated GR-at-calendar-date was compared between groups by cohort using generalized linear mixed effect models (GLMMs) with gamma distribution and log link function. We treated GR as a response variable, group as a fixed effect, and individual birth date as a random effect. Likelihood-ratio tests (LRTs) were conducted to examine the significance of groups.

The relationship between growth-rate residuals and experienced temperature was analyzed by using a linear mixed effect model (LMM). Five age classes using 18-day age intervals (age 1–18 days, age 19–36 days, age 37–54 days, age 55–72 days, and age >72 days) were included in the model as a factor variable, assuming that the relationship varies with growth stage. We treated growth-rate residual as a response variable; experienced temperature, the age classes, and the interaction between experienced temperature and the age classes as fixed effects; and individual ID as a random effect. The significance of fixed effects was examined by LRT.

All statistical tests were performed using R v.4.1.3 (R Core Team, 2022). LRTs were conducted by using the package “car” (Fox & Weisberg, 2019), and GLMMs and LMMs were conducted by using the packager “lme4” (Bates et al., 2015).

## Results

### Cohort-specific mortality rate

The vegetation index at the sampling site decreased after March due to increased water temperature (Figure 2).

**Figure 2.**
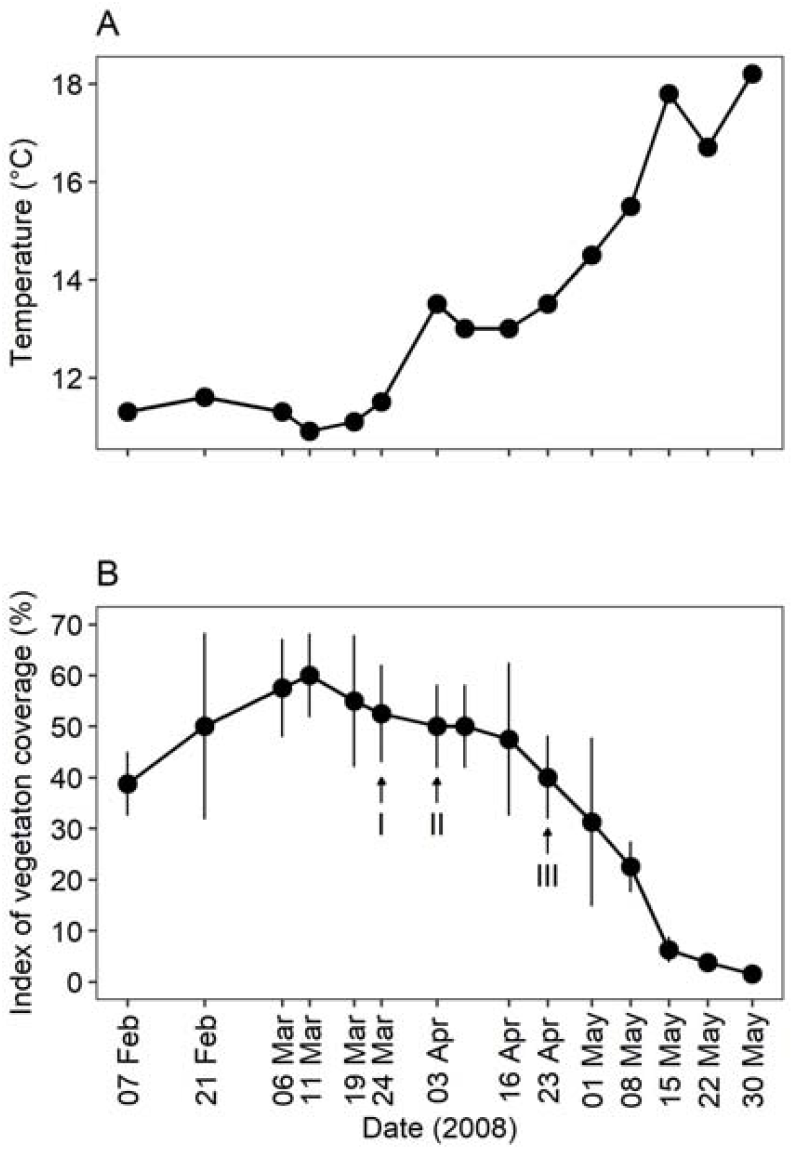
Time series of water temperature (A) and a vegetation coverage index for the macroalgal bed (B) in 2008 at the sampling site in the central Seto Inland Sea, Japan. Vertical bars denote standard deviations. Arrows denote the day of maximum mean abundance of the three cohorts. Dataset is from Kamimura & Shoji (2013).

A total of 6036 juvenile *S. cheni* were collected during the sampling period. Juvenile abundance (mean ± standard deviation; SD) was highest on 24 March (450.6 ± 327.0 inds. 100 m^−2^) and decreased up to 15 May (20.1 ± 19.0 inds. 100 m^−1^) (Figure S1). The estimated mortality coefficients (cohort I: *Z* = 0.053, *P* < 0.001, intercept = 5.45, *P* < 0.001; II: *Z* = 0.044, *P* < 0.001, intercept = 3.95, *P* < 0.001; III: *Z* = 0.071, *P* < 0.001, intercept = 2.89, *P* < 0.001) showed that the latest cohort (III) had daily mortality rates 1.8–2.7% higher than the earlier cohorts (I and II) (Figure 3 and Table 1), which is almost the same as the trend in the previous study. We processed from 14 to 33 fish from each cohort for intra- and inter-cohort comparisons of growth trajectories (Table 1). The mean TL (SD, mm) of OP and SV juvenile *S. cheni* were 25.8 (1.4) and 29.3 (2.2) for cohort I, 27.0 (1.6) and 32.0 (1.6) for cohort II, and 28.8 (1.7) and 32.8 (2.1) for cohort III, respectively.

**Figure 3.**
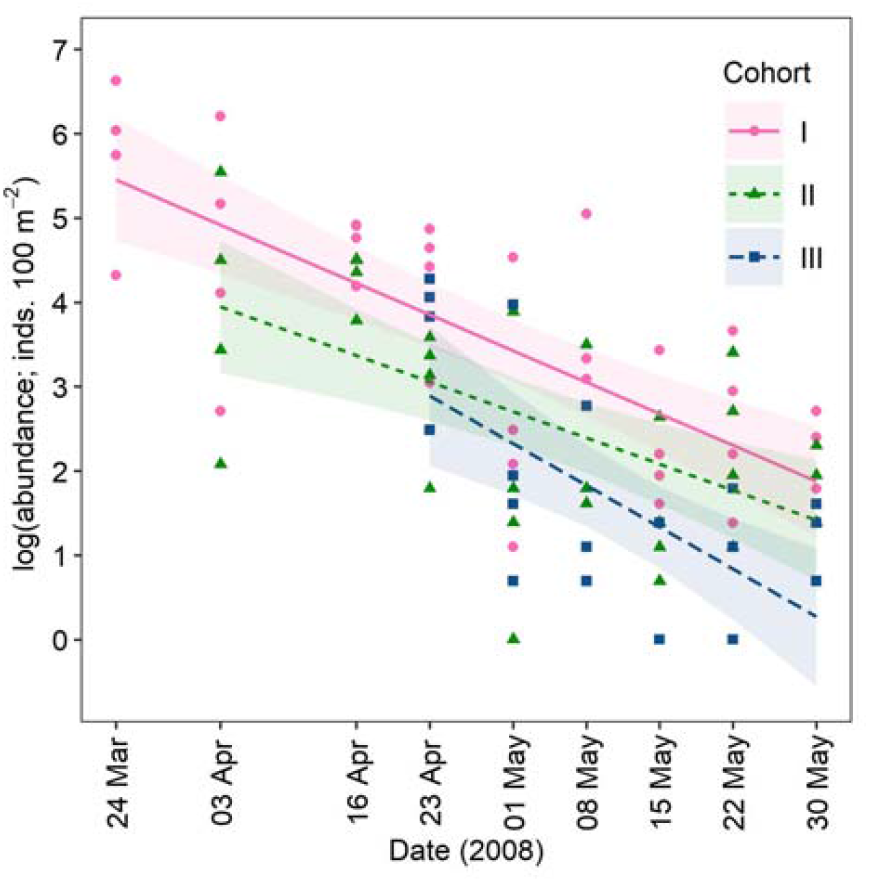
Cohort-specific changes in the natural logarithm of abundance (the number of individuals per 100 square meters; inds. 100 m^−2^) of juvenile *Sebastes cheni* by sampling date in 2008 in a macroalgal bed in the central Seto Inland Sea, Japan. Dots indicate abundance per tow by cohort. Lines and shaded areas represent linear functions and 95% confidence intervals, respectively. General linear models were fitted to the decrease in abundance to estimate the daily instantaneous mortality coefficient by cohort.

### Intra-cohort growth comparison

Mean back-calculated TL-at-age by cohort showed similar trends in OP and SV (Figure 4A). There were no significant differences between the back-calculated TL-at-age for OP or SV in the three cohorts (MANOVA; cohort I: Pillai’s Trace = 0.76, *P* = 0.34; II: Pillai’s Trace = 0.89, *P* = 0.43; III: Pillai’s Trace = 0.93, *P* = 0.41). Mean back-calculated TL-at-calendar-date by cohort also showed similar trends in OP and SV (Figure 5A). There were no significant differences between the back-calculated TLs of OP and SV in the three cohorts (MANCOVA; cohort I: Pillai’s Trace = 0.12, *P* = 0.87; II: Pillai’s Trace = 0.39, *P* = 0.29; III: Pillai’s Trace = 0.28, *P* = 0.81), whereas birth date was significantly correlated with the back-calculated TL in cohorts I and III (MANCOVA; cohort I: Pillai’s Trace = 0.39, *P* = 0.01; II: Pillai’s Trace = 0.47, *P* = 0.12; III: Pillai’s Trace = 0.82, *P* < 0.001).

**Figure 4.**
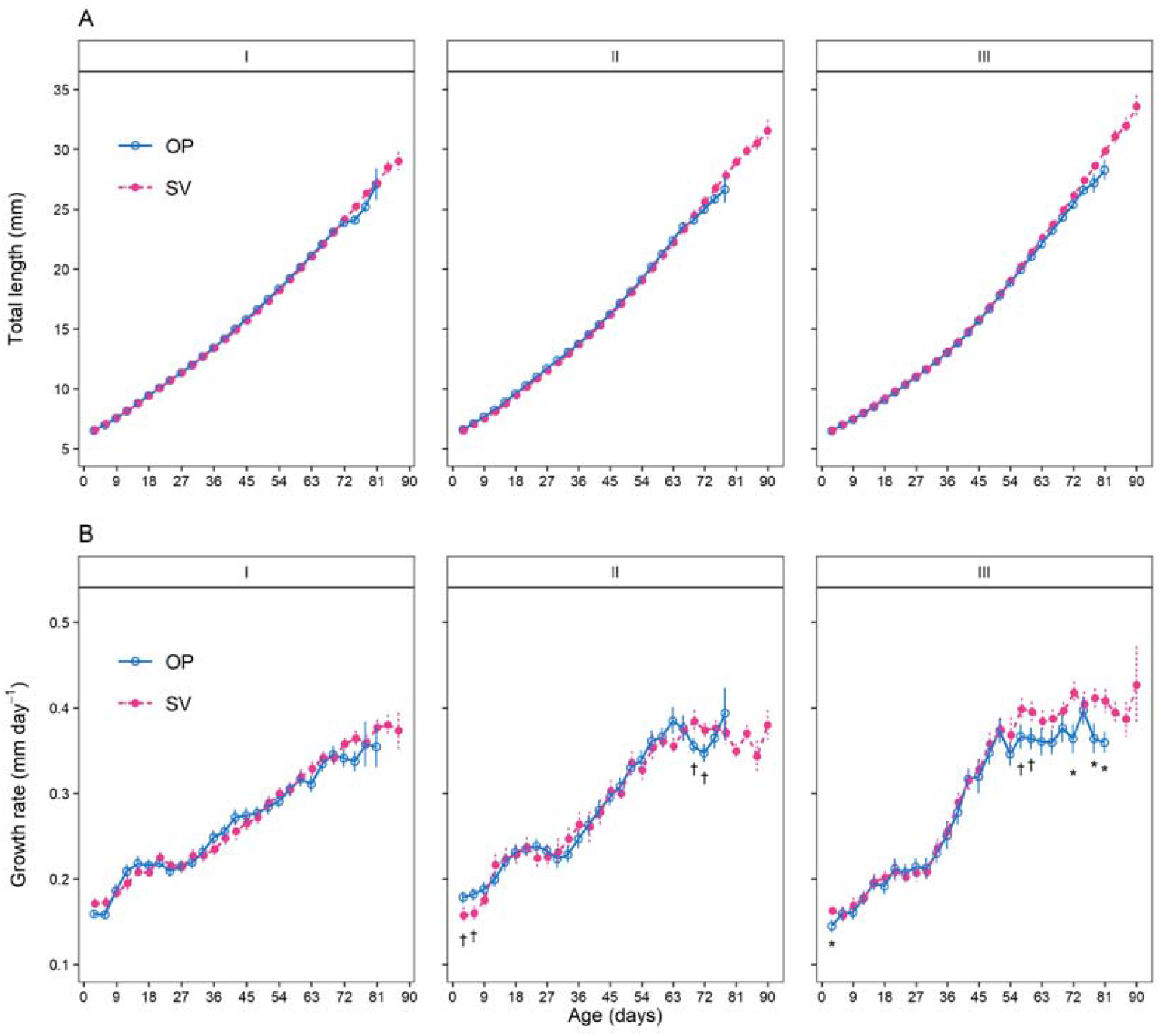
Intra-cohort age-based comparison of back-calculated mean total length (A) and growth rate (B) of *Sebastes cheni*. Open circles and solid lines denote the original population (OP) and filled circles and dotted lines denote survivors (SV). Vertical bars indicate standard errors. ^†^*P* < 0.1; \**P* < 0.05, for the comparison between OP and SV at each age.

**Figure 5.**
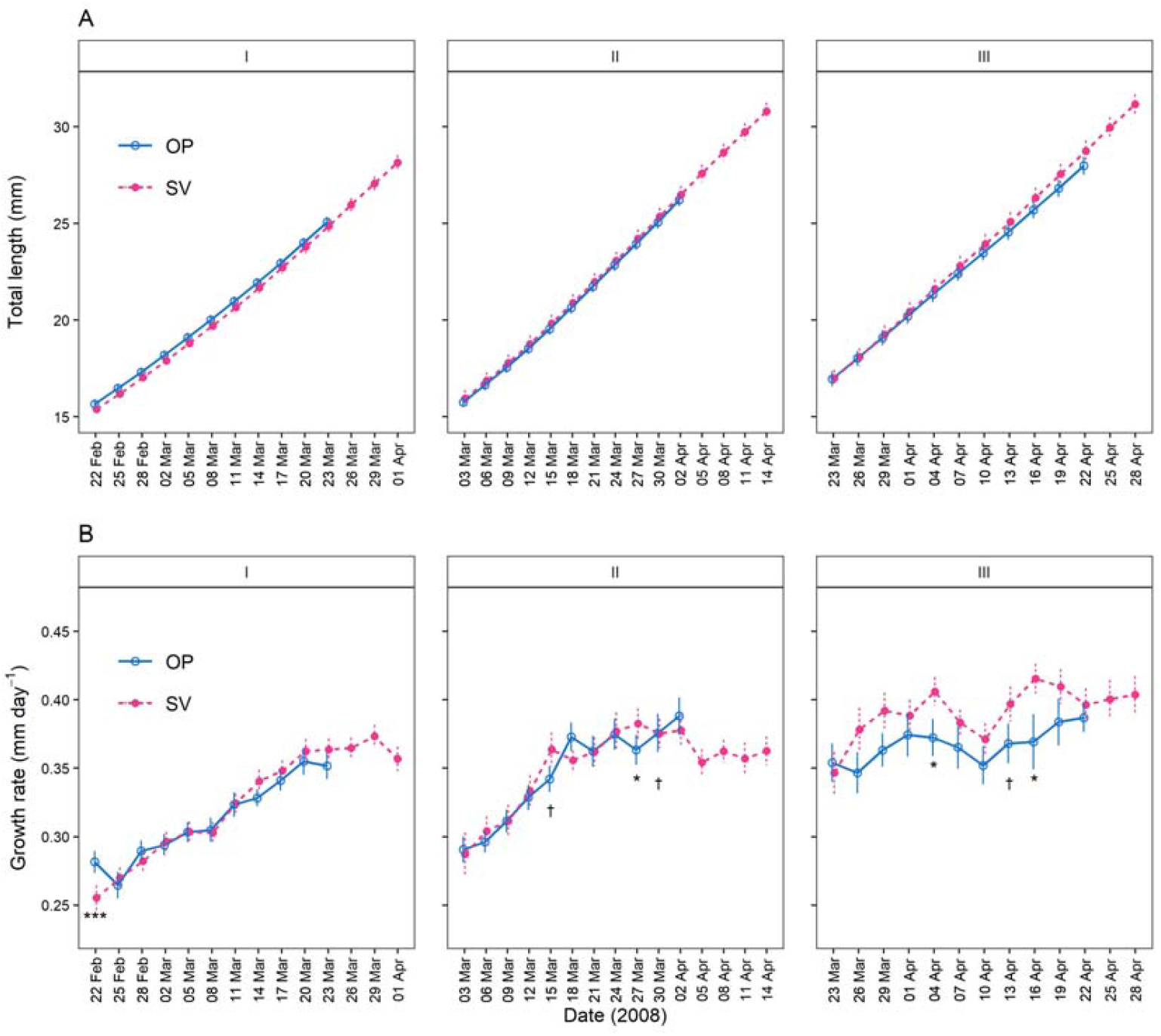
Intra-cohort date-based comparison of back-calculated mean total length (A) and growth rate (B) of *Sebastes cheni* in 2008. Open circles and solid lines denote the original population (OP) and filled circles and dotted lines denote survivors (SV). Vertical bars indicate standard errors. ^†^*P* < 0.1; \**P* < 0.05; \*\*\**P* < 0.001, for the comparison between OP and SV on each date.

Mean back-calculated GR-at-age increased up to about age 50 days (Figure 4B). In cohorts I and II, the GR of OP and SV showed similar trends, whereas there were some marginal significant differences in cohort II (Figure 4B and Table S1). In cohort III, the GR of OP and SV showed similar trends until age 54 days, except for the GR at 1–3 days (Figure 4B); after 54 days the GR was higher in SV. There were significant differences between OP and SV in the GR at age 70–72, 76–78, and 79–81 days (LRT: *P* < 0.05) and marginally significant differences in the GR at age 55–57 and 58–60 days (LRT: *P* < 0.1) (Figure 4B and Table S1). This tendency toward higher growth of SV represented the post-settlement growth–survival process because the mean back-calculated TL at age 55–57 days for cohort III was around 20 mm (Figure 4A), which corresponds to the settlement size. There were similar trends in mean back-calculated GR-at-calendar-date between OP and SV in cohorts I and II, although there were significant differences on some dates (Figure 5B and Table S2). The GR in cohort III trended higher in SV after 26 March (Figure 5B). There were significant differences in the GR between OP and SV on 1–3 April and 14–16 April (LRT: *P* < 0.05) and marginally significant differences in the GR on 11–13 April (LRT: *P* < 0.1) (Figure 5B and Table S2).

### Inter-cohort comparison of growth and experienced temperature

A logistic growth model was selected on the basis of the AIC as an appropriate model for the relationship between GR and age: linear function, GR = (2.96 × 10^−3^)age + 0.15, AIC = −10279.3; quadratic function, GR = (3.33 × 10^−3^)age – (4.33 × 10^−6^)age^2^ + 0.15, AIC = −10283.7; von Bertalanffy growth function, GR = 1.44{1 – exp[−2.56 × 10^−3^ (age + 42.7)]}, AIC = −10282.9; logistic growth function, GR = 0.50/{1 + exp[(−2.56 × 10^−2^)age + 0.81]}, AIC = −10310.8 (Figure S4).

Mean growth-rate residuals-at-age showed different trends among cohorts (Figure 6A). The mean growth-rate residual of cohort I was relatively higher for the first 20 days, after which it became lower. Conversely, that of cohort III showed a lower trend during the first 30 days, after which it was higher. The mean residual of cohort II was relatively higher for the first 30 days, after which it showed an intermediate trend.

**Figure 6.**
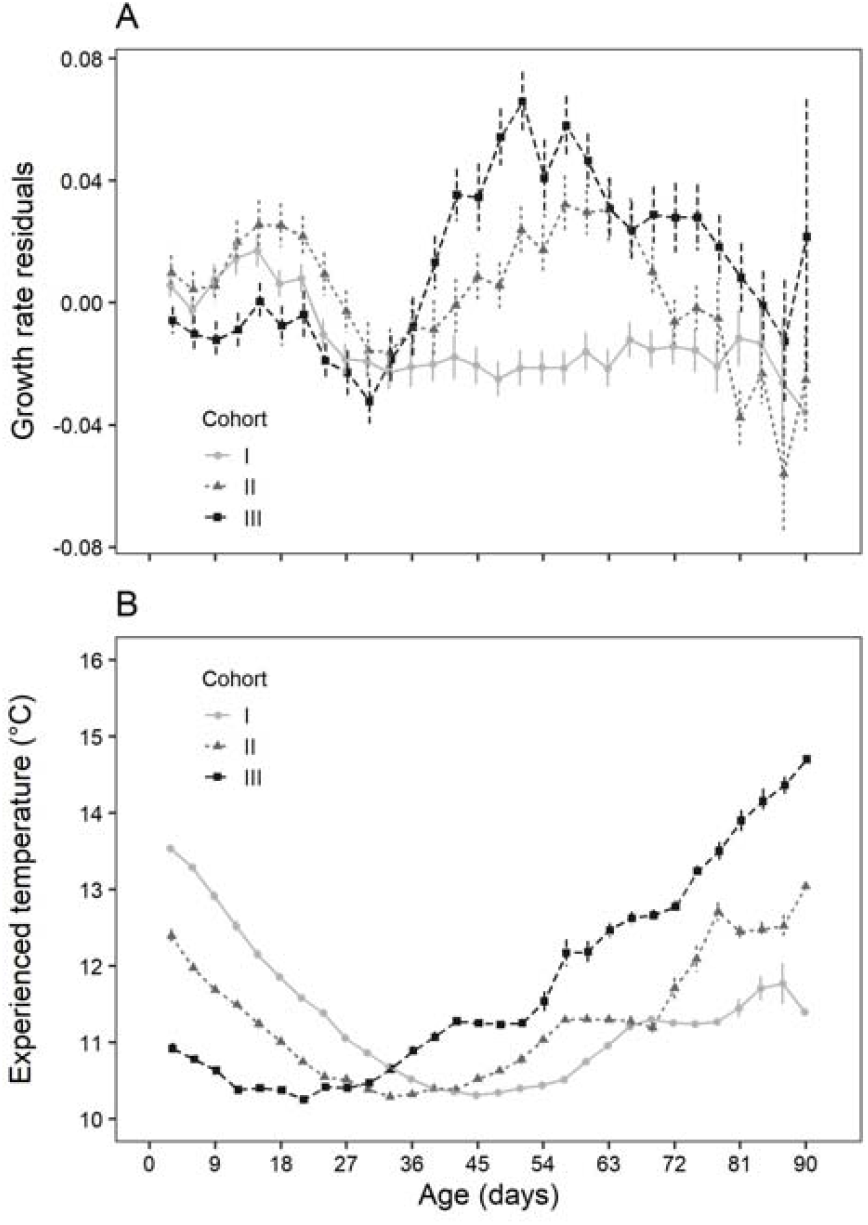
Inter-cohort comparison of back-calculated 3-day mean growth-rate residual (A) and experienced temperature (B) of *Sebastes cheni*. Circles and solid lines, triangles and dotted lines, and squares and dashed lines show cohorts I, II, and III, respectively. Vertical bars denote standard errors.

Mean experienced temperature fluctuated between 10.3 and 14.7 °C (Figure 6B). Reflecting the seasonal changes in water temperature, mean experienced temperature for cohorts I, II, and III decreased until day 45, day 33, and day 21, respectively, and then increased.

The analysis of the relationship between growth-rate residuals and experienced temperature showed that all explanatory variables were significant (LRT, experienced temperature: _χ_^2^ = 24.6, *P* < 0.001; age class: _χ_^2^ = 89.6, *P* < 0.001; experienced temperature × age class: _χ_^2^ = 102.0, *P* < 0.001). Growth-rate residuals were positively correlated with experienced temperature in all age classes except for age >72 days (Figure 7); the estimated slope of age 37–54 days was the sharpest, followed by age 19–36 and age 55–72 days. The slope of age 1–18 days was moderate.

**Figure 7.**
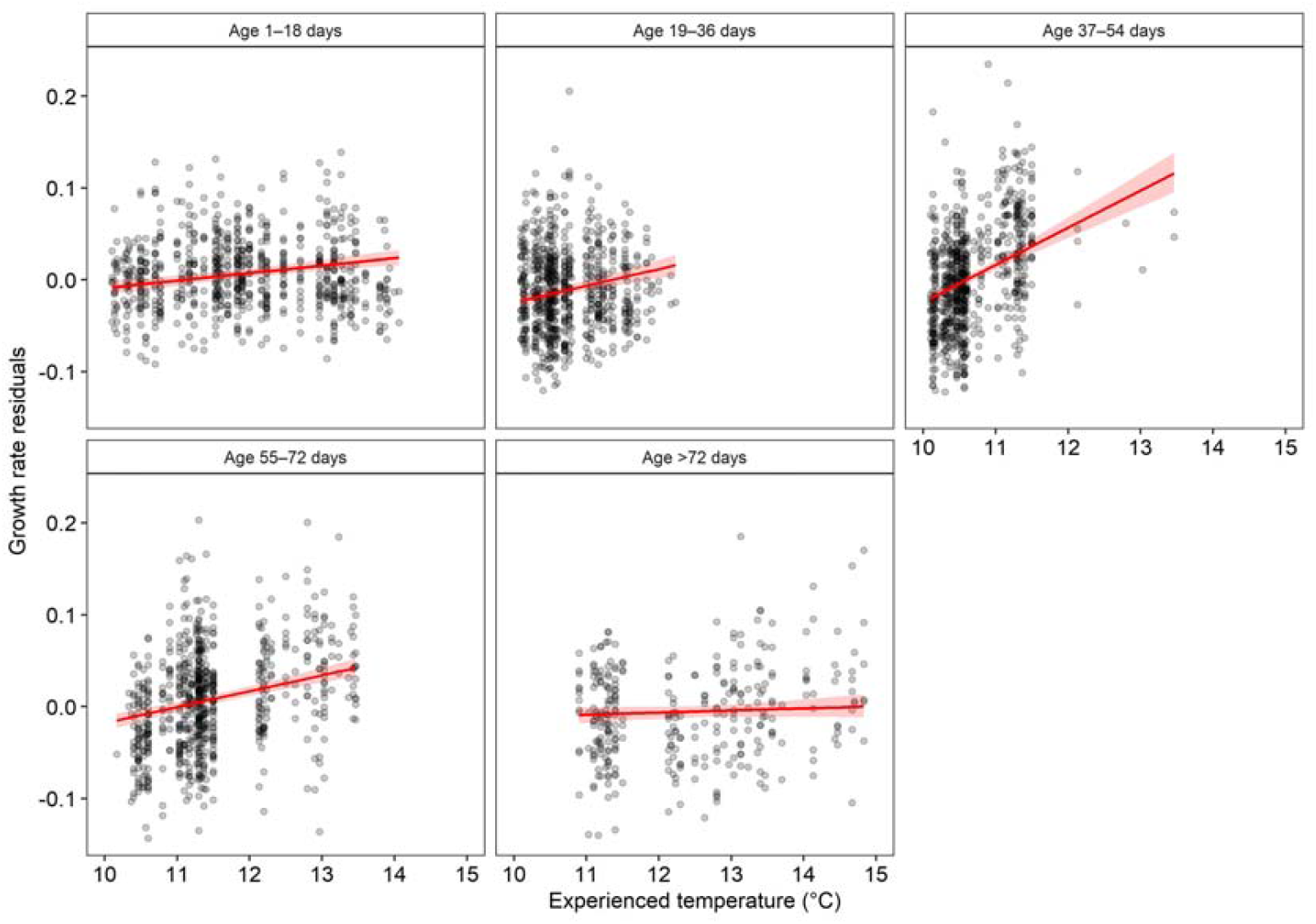
Relationship between 3-day mean growth-rate residual and experienced temperature of *Sebastes cheni* by five age-classes. Lines and shaded areas represent linear functions and 95% confidence intervals, respectively, estimated by using a general linear mixed model.

## Discussion

### Disentangling the mechanisms behind the relationship between habitat complexity and mortality of post-settlement S. cheni juveniles in a macroalgal bed

We found evidence of strong selection for faster-growing *S. cheni* juveniles—i.e., negative growth-selective mortality—in a cohort with high mortality during the post-settlement period. Repeated sampling of fish at 8-to 13-day intervals showed higher growth rates during the month before the sampling date for SV in cohort III (Figure 5B), which experienced higher mortality than cohorts I and II. In addition, the age-based comparison showed higher growth of SV for cohort III during the post-settlement period (Figure 4B). The mean back-calculated TL for the three days prior to the first sampling date for SV in cohort III (mean ± standard error, 28.74 ± 0.53 mm) was slightly greater than that of OP (27.98 ± 0.45 mm) (Figure 5A), although there were no significant differences. Hence, the juveniles seem to have experienced negative growth-selective mortality within the high-mortality cohort during the post-settlement period. In general, higher growth rate itself is positively related to burst swimming speed (Nakamura et al., 2022). Therefore, *S. cheni* growth rate during the post-settlement period is probably an indicator of anti-predator abilities, such as swimming performance, in the macroalgal bed.

A seasonal change in the mortality rate of *S. cheni* during the post-settlement period has been reported for juveniles, with higher mortality in later cohorts due to a decrease in vegetation coverage (Figure 2) (Kamimura & Shoji, 2013). The earlier cohort of juvenile *S. cheni* that settled during a period of higher vegetation coverage could probably utilize the macroalgae more effectively as shelter than the later cohorts. A day–night comparison of fish assemblages in the macroalgal bed showed an increase in the biomass of piscivorous fishes such as adult rockfish *Sebastes* spp. and conger eel *Conger myriaster* during the nighttime (Kinoshita et al., 2014), confirmed to be nocturnal visitors by acoustic telemetry (Shoji et al., 2017). In addition, stomach contents analysis of these fishes has shown that they are likely to be dominant predators of juvenile *S. cheni* (Kinoshita et al., 2014). However, there were no significant seasonal changes in predator biomass from March through May (Kinoshita et al., 2014). Therefore, it is plausible that the susceptibility of juvenile *S. cheni* to predation was affected by the seasonal change in vegetation coverage under similar levels of predator biomass.

Furthermore, differences in mortality rate among cohorts show that post-settlement juveniles experience negative size-selective mortality under lower vegetation coverage. In other words, the fact that the three cohorts co-inhabited the macroalgal bed after 23 April, and that only cohort III, which consisted of the youngest and smallest juveniles among the three cohorts, underwent higher mortality (Figure 3) should be a reflection of the consequences of negative size-selective mortality. This suggestion is supported by the following results. The mean TL of OP on 23 April of cohort III (mean ± SD; 28.8 ± 1.7 mm) (Table 1) was in fact smaller than mean TLs on that date of cohorts I (40.6 ± 1.7 mm) and II (34.1 ± 1.0 mm), calculated using the dataset from a previous study (Kamimura & Shoji, 2013). Moreover, even when the period for estimating mortality for all cohorts was aligned after 3 April or after 24 April, the results remained largely the same as the original estimate, that is, cohort III had the highest mortality rate among the three cohorts (Figure S5). This observation also suggests that if juvenile body size is greater than the limit for high predation risk (around 30 mm TL), mortality can be reduced even under low vegetation.

In summary, the intra- and inter-cohort comparisons for mortality rates and growth trajectories suggest that lower vegetation coverage on the macroalgal bed increased the daily mortality rate of juvenile *S. cheni* by 1.8–2.7% (Figure 3) through both negative size- and growth-selective predation. Although previous studies have demonstrated that higher habitat complexity in vegetated habitats decreases larval and juvenile mortality rates (e.g., Horinouchi, 2007; Johnson, 2006), to our knowledge, no study has shown that it influences the growth–survival relationship in early growth stages of fish.

Such results suggest that vegetation coverage in macroalgal beds plays a critical role in controlling the relationship between predators and growth of juvenile *S. cheni*. Therefore, vegetation coverage (habitat complexity) should be considered as a factor affecting growth–survival mechanisms in fish early life stages in addition to two other key factors: the predator field (temporal changes in predator abundance and composition encountered by larvae) and the prey field (absolute growth rate of larvae and juveniles and its variability for each cohort) (Takasuka et al., 2017). In general, the bigger-is-better and growth-selective predation hypotheses conceive a monotonic decrease in mortality rate with increases in somatic size or growth rate (e.g., Leggett & Frank, 2008). Here, we add a new attribute, habitat complexity, to that conception as a regulatory function (Figure 8). It states that although juveniles in less structured habitats suffer higher mortality than juveniles in highly structured habitats through a combination of negative size- and growth-selective predation, increases in somatic size rapidly reduce mortality. The mechanisms behind size-selective mortality, growth-selective mortality, and the stage-duration are interconnected, or interact, and tend to depend on species, populations, or seasonal cohorts (Takasuka et al., 2017). Our results can help to understand the dynamics of survival mechanisms in fish early life stages. On the other hand, a recent study suggested that connectivity in the seascape between nearshore habitats (seagrass, kelp forest, and sand habitats) also influences post-settlement juvenile density, growth, and survival in nursery habitats (Olson et al., 2019). The site of the present study, Abajima Island, is isolated from other islands and surrounded by waters with depths >50 m, which prevents juveniles from emigrating from the study site. It is therefore plausible that *S. cheni* juveniles did not actively migrate out of the macroalgal bed during the survey period (Kamimura & Shoji, 2013). Future macro-scale studies considering habitat connectivity would help to clarify survival processes among substrate-associated larvae and juveniles and to evaluate the nursery function of vegetated habitats.

**Figure 8.**
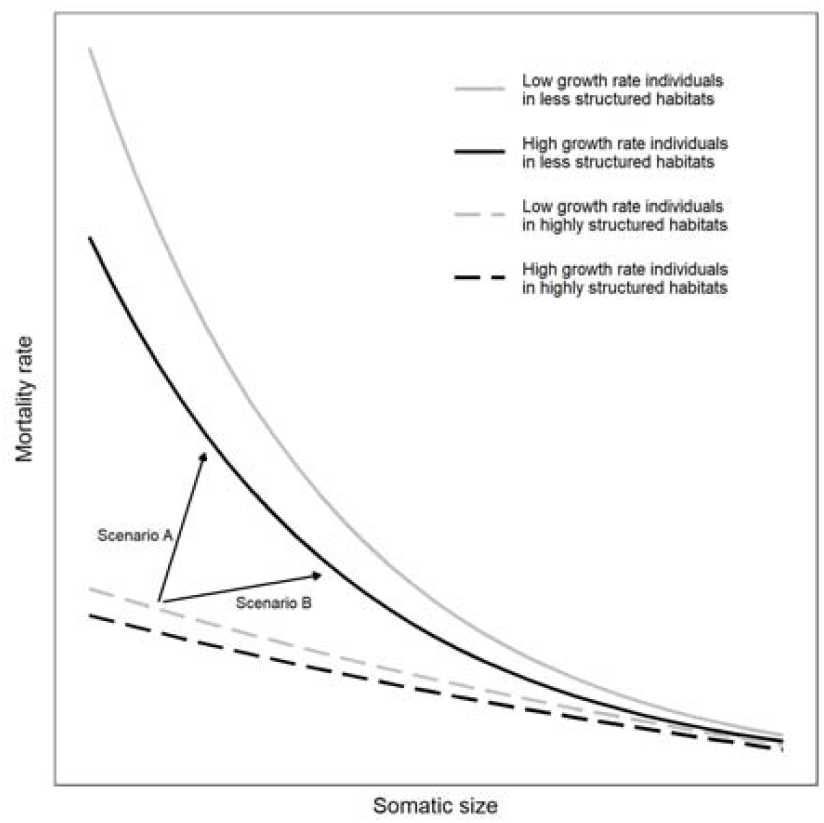
Conceptual diagram of the relationship between somatic size and mortality rate as regulated by habitat complexity, such as vegetation coverage. Mortality rate is lower in highly structured habitats than in less structured habitats. In less structured habitats, individuals with low growth rate tend to suffer higher mortality than those with a higher growth rate within a population consisting of fish of the same size. Arrows depict the seasonal changes in relative mortality rates from a sharp decrease in habitat complexity accompanying a rapid increase in temperature (Scenario A) or from a slow decrease in habitat complexity accompanying a gradual increase in temperature (Scenario B).

Kinoshita et al. (2014) compared back-calculated growth of juvenile *S. cheni* survivors with that of those ingested by predators. Their results provide evidence for negative size-selective predation on juveniles, but not for growth-selective predation, which we demonstrated in the present study. This difference is probably due to differences in survey design. The previous study aggregated individual growth data from seven sampling dates between mid-March and late May, when vegetation coverage is likely to change dramatically. Therefore, their use of merged growth trajectories for juveniles that experienced different conditions of vegetation coverage likely prevented the detection of any negative growth-selective predation.

### Growth differences among cohorts and temperature effects on growth and survival

The early growth histories of *S. cheni* are probably affected by the consequences of morphological and physiological changes. Growth-rate residuals of *S. cheni* showed a consistent pattern, that is, an increase in growth rate after birth, a decrease in growth until around age 30 days (around 12 mm TL), and another increase after that (Figure 6A) as also reported in a previous study (Plaza et al., 2001). That study suggested that the growth pattern could be explained by ontogenetic changes, and our results are consistent with this explanation. At a TL around 12 mm, *S. cheni* is in the postflexion larval stage (Nagasawa et al., 2000), when feeding habits change and the habitat shifts from pelagic to benthic; these changes can affect metabolic osmoregulation and prey availability through reduction in temperature and light levels (Plaza et al., 2001). This is probably the common reason for the growth stagnation of *S. cheni* around age 30 days in all cohorts (Figure 6A) and the weak positive relationship between temperature and growth until age 40 days (Figure 7).

The relationship between water temperature and *S. cheni* growth combined with the seasonal trend in temperature suggests that experienced temperature largely determines the growth pattern of *S. cheni*. Water temperature positively affected *S. cheni* growth between 10 and 15 °C, although the strength of the effects differed among age classes, peaking at age 37–54 days (14–19 mm TL) (Figure 7), the pre-settlement period. A previous study also reported that higher sea-surface temperature enhanced the pre-settlement growth of *S. melanops, S. caurinus, S. maliger*, and *S. auriculatus* (Markel & Shurin, 2020). Therefore, the latest cohort in our study experienced a consistent temperature increase and exhibited rapid growth in the pre-settlement stage.

The reason why the temperature effects on growth weakened after settlement (age 55–57 days) can be explained by the biological characteristics of *S. cheni*. Because post-settlement *Sebastes* juveniles are probably exposed to stronger intra-specific competition for food as they exhibit schooling and aggregating behavior in shallow waters, food availability is likely to be more important for *S. cheni* growth than temperature. In fact, a previous study suggested a possible negative density-dependence on growth in *Sebastes* juveniles during the post-settlement period (Plaza et al., 2002). Further studies of consumption rates and food availability for post-settlement *S. cheni* juveniles would help to understand their density-dependence in growth.

A more gradual temperature increase in the macroalgal bed seems to be more favorable for the survival of post-settlement *S. cheni*, because the increase in water temperature experienced by the later cohort resulted in higher mortality through a decrease in vegetation coverage (Figure 3), whereas there was no rapid acceleration in growth (Figure 7). In other words, increases in growth rate induced by rapid seasonal increases in water temperature likely do not fully balance the increases in mortality rate. Therefore, the speed of the transition from high to low vegetation coverage may be important in determining the total mortality of an *S. cheni* population during the post-settlement period. If there is a rapid decline in vegetation coverage due to a rapid temperature increase, the population mortality rate is higher (scenario A in Figure 8) than if the decline is slower (scenario B in Figure 8). The lower survival index in later cohorts estimated in a previous study (Kamimura & Shoji, 2013) also supports this suggestion. In addition, Terazono et al. (2012) suggested that increased temperatures caused by global warming will shorten the vegetative period of temperate *Sargassum* in spring and this may have a negative impact on local fish populations. The effects of higher temperatures on survival of larvae and juveniles that are dependent on vegetated habitats require further investigation.

## Conclusions

Our study confirmed that vegetation coverage in a macroalgal bed was critical for the quantity and quality of juvenile *S. cheni* surviving during the post-settlement period. The probability of survival for juvenile *S. cheni* that settled earlier in the season under lower water temperatures was higher because of higher vegetation coverage (i.e., more effective shelter). By contrast, the later cohort that settled during a time of low vegetation coverage had lower survival due to strong negative size- and growth-selective predation by nighttime predators, that is, bigger and faster-growing fish have a better chance of survival under low vegetation coverage because they have better physiological responses (such as burst swimming speed). The relationship between experienced temperature and growth rate for *S. cheni* showed that higher water temperature may have resulted in a trend toward higher growth in the later cohort. However, the magnitude of the temperature effects on growth seems to have weakened during the post-settlement stage as a result of the effects of negative density-dependent growth. Therefore, we suggest that a gradual decrease in vegetation coverage with a slower increase in temperature might be ideal for *S. cheni* survival during the post-settlement period. Our findings should be useful for management of this fish species and for conservation of vegetated habitats under the increasing effects of global warming, and suggests the necessity of further studies into the effects of increasing temperature on larval and juvenile survival of species dependent on temperate algae.

## Acknowledgements

We appreciate S. Iwasaki, Y. Iwamoto, T. Morita, K. Hirai, S. Inoue, K. Mishiro, K. Mizuno, H. Kinoshita, S. Toshito, and K. Mohri, Takehara Fisheries Research Station, Hiroshima University, for their help with surveys. We thank K. Hirakawa for providing data for daily water temperature. The authors also thank English-speaking professional editors from ELSS, Inc., for English-language proofreading. This work was supported by the Japan Society for the Promotion of Science (JSPS) KAKENHI Grant Number JP10J08287.

## Author contributions

YK and JS conceived the ideas and collected the data. YK performed the data analysis and drafted the original manuscript. YK and JS reviewed and revised the draft manuscript and approved the final version for submission.

## Data availability statement

The data that support the findings of this study are available from the corresponding author, YK, upon reasonable request.

## Supporting information

Additional supporting information may be found in the online version of the article at the publisher’s website.

