## Supporting Information for "Highly structured habitats mitigate size- and growth-selective mortality of post-settlement juvenile fish"

Table S1. Results of generalized linear models with likelihood-ratio tests for the age-based comparison of 3-day mean back-calculated growth rates between groups (OP, original population; SV, survivors) of juvenile *Sebastes cheni* from the central Seto Inland Sea, Japan. ^†^*P* < 0.1; **P* < 0.05.

| Cohort | Age (days) | Sample size | |  | Likelihood-ratio test | |
| --- | --- | --- | --- | --- | --- | --- |
|  |  | OP | SV |  | *χ*^2^ value | *P* value |
| I | 1–3 | 25 | 33 |  | 2.446 | 0.118 |
|  | 4–6 | 25 | 33 |  | 2.199 | 0.138 |
|  | 7–9 | 25 | 33 |  | 0.049 | 0.825 |
|  | 10–12 | 25 | 33 |  | 1.461 | 0.227 |
|  | 13–15 | 25 | 33 |  | 0.817 | 0.366 |
|  | 16–18 | 25 | 33 |  | 0.857 | 0.355 |
|  | 19–21 | 25 | 33 |  | 0.483 | 0.487 |
|  | 22–24 | 25 | 33 |  | 0.436 | 0.509 |
|  | 25–27 | 25 | 33 |  | 0.002 | 0.969 |
|  | 28–30 | 25 | 33 |  | 0.524 | 0.469 |
|  | 31–33 | 25 | 33 |  | 0.128 | 0.720 |
|  | 34–36 | 25 | 33 |  | 0.988 | 0.320 |
|  | 37–39 | 25 | 33 |  | 0.425 | 0.515 |
|  | 40–42 | 25 | 33 |  | 1.101 | 0.294 |
|  | 43–45 | 25 | 33 |  | 0.462 | 0.497 |
|  | 46–48 | 25 | 33 |  | 0.211 | 0.646 |
|  | 49–51 | 25 | 33 |  | 0.155 | 0.694 |
|  | 52–54 | 25 | 33 |  | 0.572 | 0.449 |
|  | 55–57 | 25 | 33 |  | 0.001 | 0.976 |
|  | 58–60 | 25 | 33 |  | 0.032 | 0.858 |
|  | 61–63 | 25 | 33 |  | 2.136 | 0.144 |
|  | 64–66 | 25 | 33 |  | 0.356 | 0.551 |
|  | 67–69 | 25 | 33 |  | 0.112 | 0.738 |
|  | 70–72 | 20 | 33 |  | 1.836 | 0.175 |
|  | 73–75 | 11 | 33 |  | 2.786 | 0.095 |
|  | 76–78 | 6 | 33 |  | 0.007 | 0.934 |
|  | 79–81 | 2 | 25 |  | 0.423 | 0.516 |
| II | 1–3 | 21 | 14 |  | 3.349 | 0.067 ^†^ |
|  | 4–6 | 21 | 14 |  | 3.723 | 0.054 ^†^ |
|  | 7–9 | 21 | 14 |  | 1.413 | 0.235 |
|  | 10–12 | 21 | 14 |  | 1.417 | 0.234 |
|  | 13–15 | 21 | 14 |  | 0.053 | 0.818 |
|  | 16–18 | 21 | 14 |  | 0.025 | 0.875 |
|  | 19–21 | 21 | 14 |  | 0.021 | 0.884 |
|  | 22–24 | 21 | 14 |  | 0.976 | 0.323 |
|  | 25–27 | 21 | 14 |  | 0.250 | 0.617 |
|  | 28–30 | 21 | 14 |  | 0.158 | 0.691 |
|  | 31–33 | 21 | 14 |  | 1.318 | 0.251 |
|  | 34–36 | 21 | 14 |  | 0.860 | 0.354 |
|  | 37–39 | 21 | 14 |  | 0.013 | 0.910 |
|  | 40–42 | 21 | 14 |  | 0.021 | 0.886 |
|  | 43–45 | 21 | 14 |  | 0.211 | 0.646 |
|  | 46–48 | 21 | 14 |  | 0.273 | 0.601 |
|  | 49–51 | 21 | 14 |  | 0.116 | 0.733 |
|  | 52–54 | 21 | 14 |  | 0.535 | 0.465 |
|  | 55–57 | 21 | 14 |  | 0.148 | 0.701 |
|  | 58–60 | 21 | 14 |  | 0.117 | 0.732 |
|  | 61–63 | 21 | 14 |  | 2.126 | 0.145 |
|  | 64–66 | 21 | 14 |  | 0.020 | 0.888 |
|  | 67–69 | 17 | 14 |  | 3.313 | 0.069 ^†^ |
|  | 70–72 | 16 | 14 |  | 3.423 | 0.064 ^†^ |
|  | 73–75 | 10 | 14 |  | 0.609 | 0.435 |
|  | 76–78 | 3 | 14 |  | 0.485 | 0.486 |
| III | 1–3 | 14 | 16 |  | 4.173 | 0.041 * |
|  | 4–6 | 14 | 16 |  | 0.018 | 0.892 |
|  | 7–9 | 14 | 16 |  | 0.398 | 0.528 |
|  | 10–12 | 14 | 16 |  | 0.003 | 0.959 |
|  | 13–15 | 14 | 16 |  | 0.008 | 0.929 |
|  | 16–18 | 14 | 16 |  | 0.547 | 0.460 |
|  | 19–21 | 14 | 16 |  | 0.042 | 0.837 |
|  | 22–24 | 14 | 16 |  | 0.186 | 0.666 |
|  | 25–27 | 14 | 16 |  | 0.210 | 0.647 |
|  | 28–30 | 14 | 16 |  | 0.086 | 0.769 |
|  | 31–33 | 14 | 16 |  | 0.126 | 0.723 |
|  | 34–36 | 14 | 16 |  | 0.065 | 0.798 |
|  | 37–39 | 14 | 16 |  | 0.454 | 0.501 |
|  | 40–42 | 14 | 16 |  | 0.012 | 0.913 |
|  | 43–45 | 14 | 16 |  | 0.136 | 0.712 |
|  | 46–48 | 14 | 16 |  | 0.289 | 0.591 |
|  | 49–51 | 14 | 16 |  | 0.004 | 0.948 |
|  | 52–54 | 14 | 16 |  | 0.778 | 0.378 |
|  | 55–57 | 14 | 16 |  | 2.912 | 0.088 ^†^ |
|  | 58–60 | 14 | 16 |  | 3.356 | 0.067 ^†^ |
|  | 61–63 | 14 | 16 |  | 1.361 | 0.243 |
|  | 64–66 | 14 | 16 |  | 2.235 | 0.135 |
|  | 67–69 | 14 | 16 |  | 1.197 | 0.274 |
|  | 70–72 | 13 | 16 |  | 6.416 | 0.011 * |
|  | 73–75 | 13 | 16 |  | 0.109 | 0.741 |
|  | 76–78 | 6 | 16 |  | 4.783 | 0.029 * |
|  | 79–81 | 6 | 16 |  | 4.544 | 0.033 * |

Table S2. Results of generalized linear mixed effects models with likelihood-ratio tests for the date-based comparison of 3-day mean back-calculated growth rates between groups (OP, original population; SV, survivors) of juvenile *Sebastes cheni* from the central Seto Inland Sea, Japan. ^†^*P* < 0.1; **P* < 0.05; ****P* < 0.001.

| Cohort | Calendar date (2008) | Sample size | |  | Likelihood-ratio test | |
| --- | --- | --- | --- | --- | --- | --- |
|  |  | OP | SV |  | *χ*^2^ value | *P* value |
| I | 20–22 Feb | 25 | 33 |  | 17.359 | <0.001 *** |
|  | 23–25 Feb | 25 | 33 |  | 0.158 | 0.691 |
|  | 26–28 Feb | 25 | 33 |  | 0.206 | 0.650 |
|  | 31 Jan–2 Mar | 25 | 33 |  | 1.215 | 0.270 |
|  | 3–5 Mar | 25 | 33 |  | 0.002 | 0.968 |
|  | 6–8 Mar | 25 | 33 |  | 0.067 | 0.796 |
|  | 9–11 Mar | 25 | 33 |  | 0.034 | 0.855 |
|  | 12–14 Mar | 25 | 33 |  | 1.374 | 0.241 |
|  | 15–17 Mar | 25 | 33 |  | 0.504 | 0.478 |
|  | 18–20 Mar | 25 | 33 |  | 0.316 | 0.574 |
|  | 21–23 Mar | 25 | 33 |  | 1.243 | 0.265 |
| II | 1–3 Mar | 21 | 14 |  | 0.141 | 0.707 |
|  | 4–6 Mar | 21 | 14 |  | 0.394 | 0.530 |
|  | 7–9 Mar | 21 | 14 |  | 0.160 | 0.689 |
|  | 10–12 Mar | 21 | 14 |  | 0.143 | 0.705 |
|  | 13–15 Mar | 21 | 14 |  | 3.691 | 0.055 ^†^ |
|  | 16–18 Mar | 21 | 14 |  | 1.163 | 0.281 |
|  | 19–21 Mar | 21 | 14 |  | 0.312 | 0.576 |
|  | 22–24 Mar | 21 | 14 |  | 0.361 | 0.548 |
|  | 25–27 Mar | 21 | 14 |  | 5.051 | 0.025 * |
|  | 28–30 Mar | 21 | 14 |  | 3.064 | 0.080 ^†^ |
|  | 31 Mar–2 Apr | 21 | 14 |  | 0.083 | 0.774 |
| III | 21–23 Mar | 14 | 16 |  | 0.251 | 0.616 |
|  | 24–26 Mar | 14 | 16 |  | 2.003 | 0.157 |
|  | 27–29 Mar | 14 | 16 |  | 2.689 | 0.101 |
|  | 30 Mar–1 Apr | 14 | 16 |  | 0.507 | 0.477 |
|  | 2–4 Apr | 14 | 16 |  | 3.914 | 0.048 * |
|  | 5–7 Apr | 14 | 16 |  | 0.403 | 0.526 |
|  | 8–10 Apr | 14 | 16 |  | 0.308 | 0.579 |
|  | 11–13 Apr | 14 | 16 |  | 3.491 | 0.062 ^†^ |
|  | 14–16 Apr | 14 | 16 |  | 4.192 | 0.041 * |
|  | 17–19 Apr | 14 | 16 |  | 1.129 | 0.288 |

Figure S1. Changes in mean abundance (number of individuals per 100 square meters, inds. 100 m^−2^) of juvenile *Sebastes cheni* collected in 2008 from a macroalgal bed in the central Seto Inland Sea, Japan. Grey dots show raw data and vertical bars denote standard deviations. Data are from Kamimura & Shoji (2013).

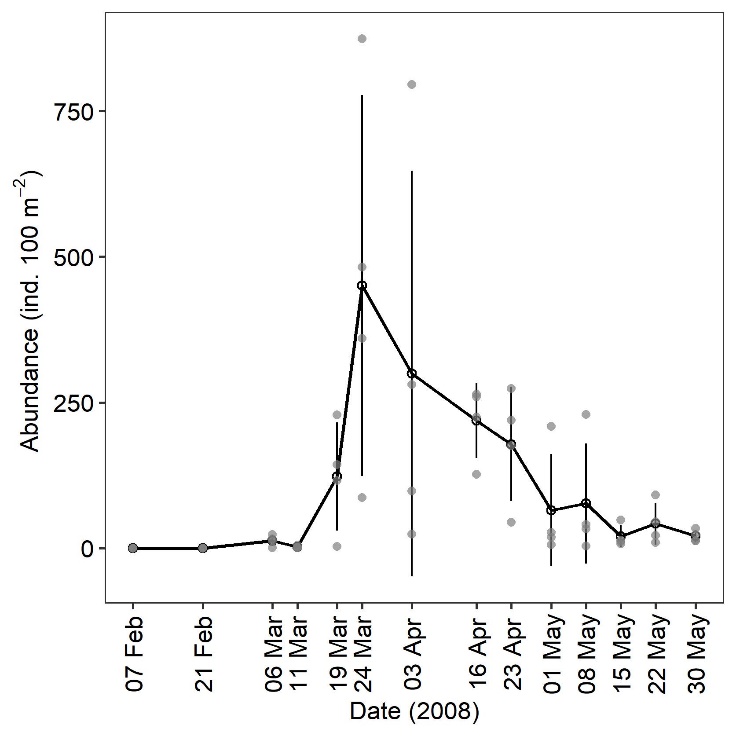

Figure S2. Daily water temperature near the sampling site in 2008 obtained from the Hiroshima Prefectural Farming Fisheries Center.

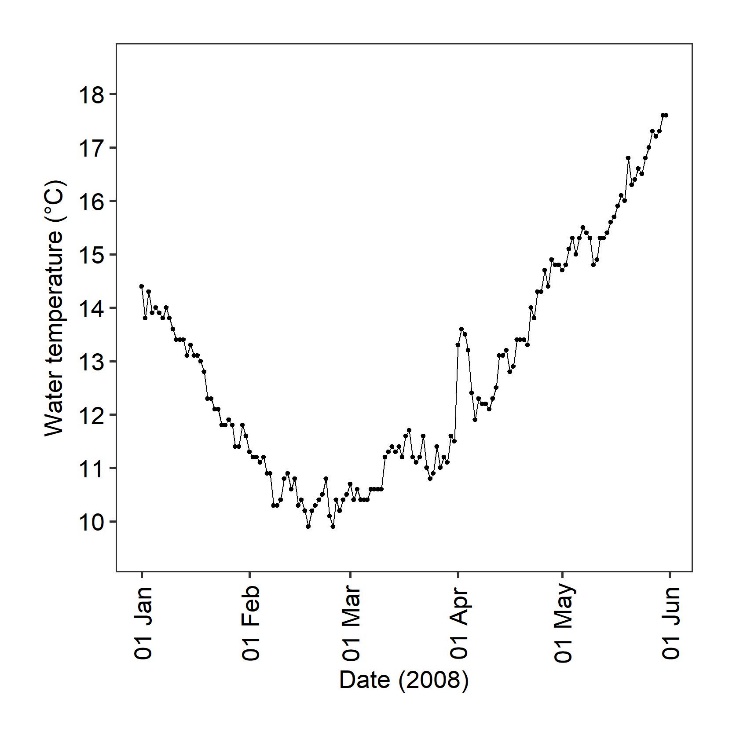

Figure S3. Birth-date distribution of *Sebastes cheni* used for estimating growth trajectories by cohort (I, II, and III). Each cohort covers a two-week birth-date period in 2008. Blue and red bars denote the original population (OP) and survivors (SV), respectively.

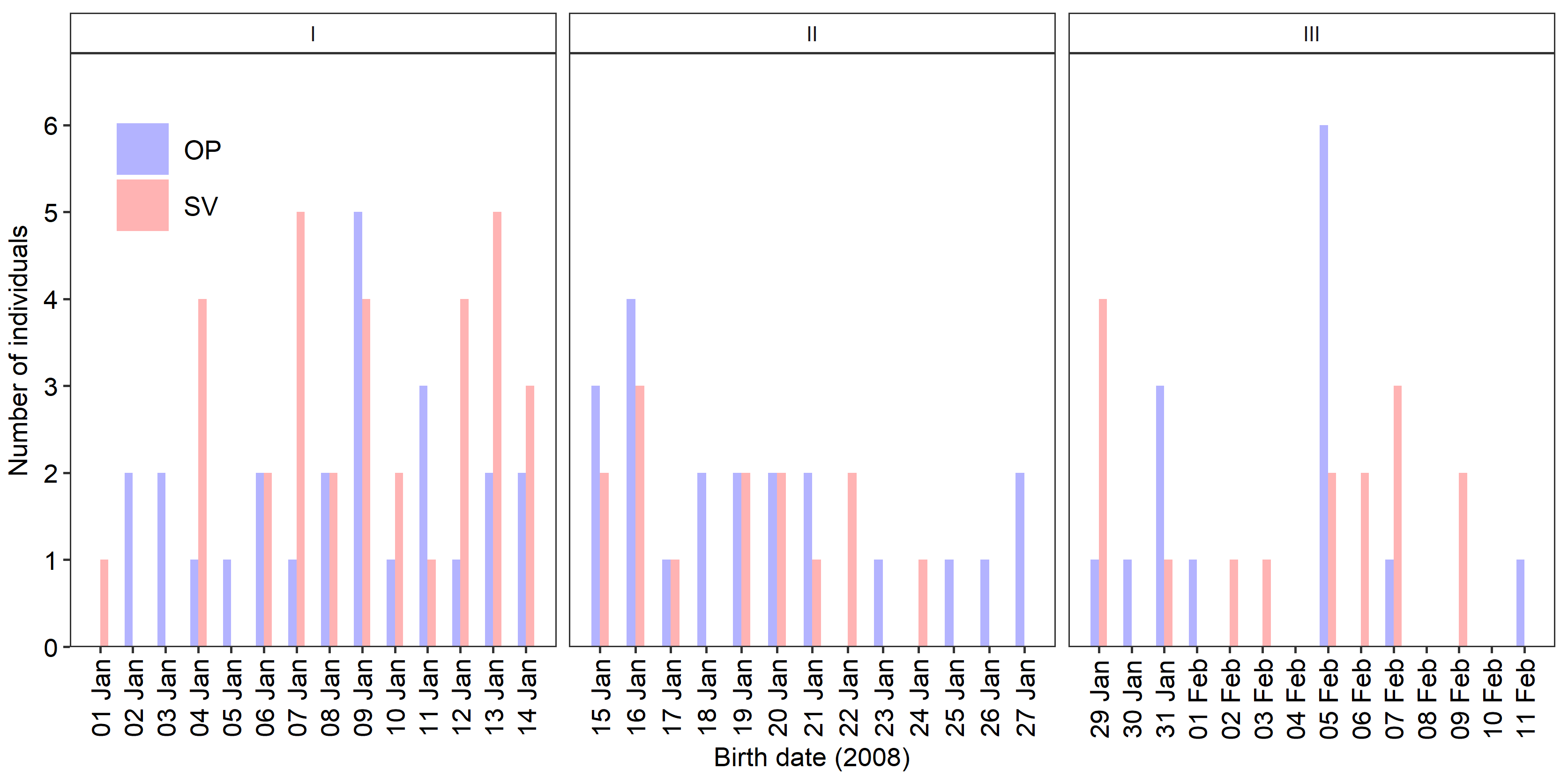

Figure S4. The relationship between 3-day mean growth rate and age for *Sebastes cheni* from a macroalgal bed in the central Seto Inland Sea, Japan. Grey dots show raw data and the black line represents the estimated values of a logistic growth model. Red shaded areas show 95% confidence intervals. The model was estimated by a generalized linear model using a Gamma distribution and log link function. Residuals of the model were used as our estimate of de-trended growth rate.

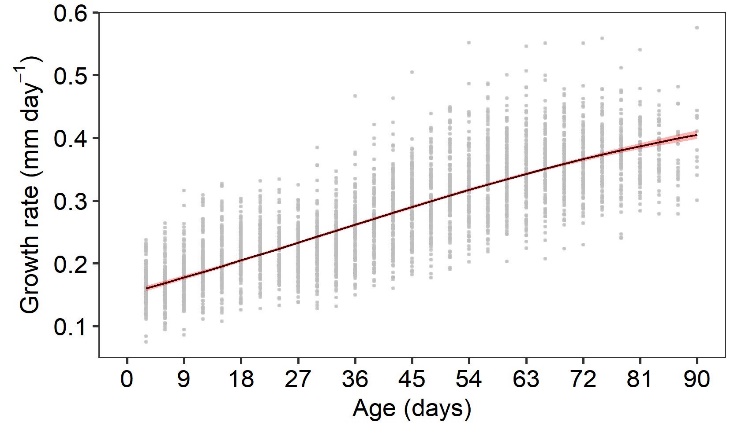

Figure S5. Cohort-specific changes in the natural logarithm of abundance (the number of individuals per 100 square meters; inds. 100 m^−2^) of juvenile *Sebastes cheni* by sampling date in a macroalgal bed in the central Seto Inland Sea, Japan, in 2008. Each cohort covers a two-week birth-date period. Dots indicate abundances per tow by cohort. Lines and shaded areas represent linear functions and 95% confidence intervals, respectively. General linear models were fitted to the decrease in abundance to estimate the daily instantaneous mortality coefficients (*Z*, day^–1^) by cohort and by the three starting dates (24 March, 3 April, and 23 April).

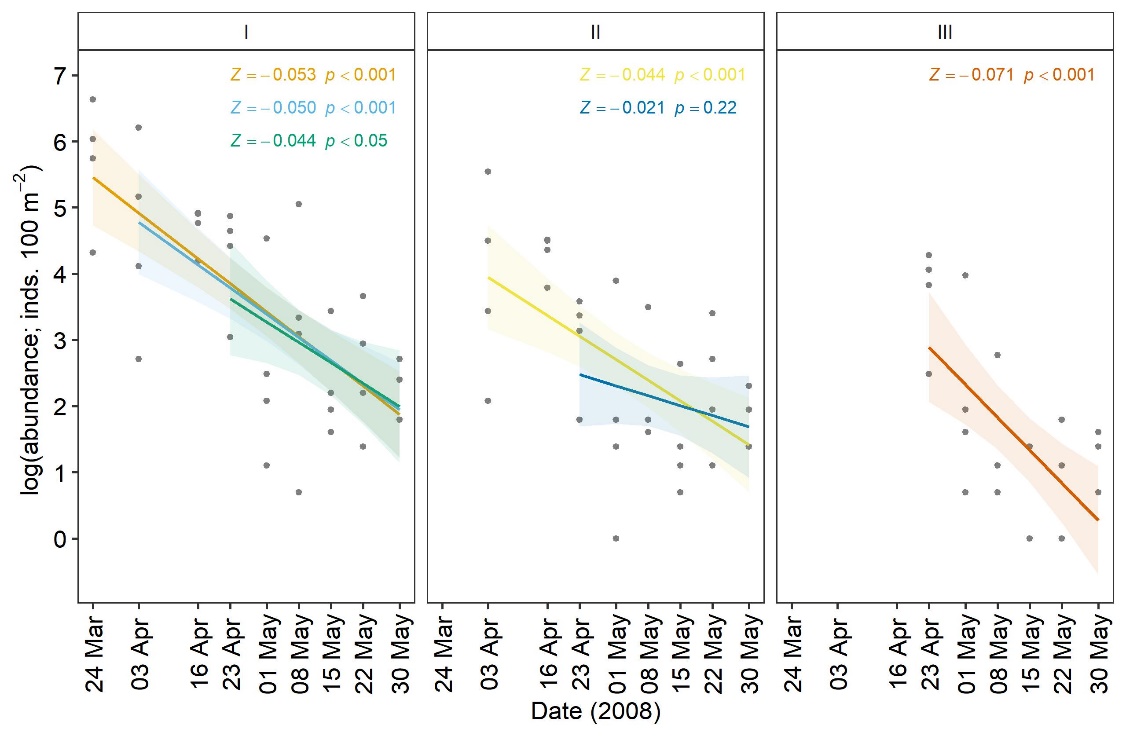
